# Non-invasive mapping of the temporal processing hierarchy in the human visual cortex

**DOI:** 10.1101/2025.05.12.653426

**Authors:** Katharina Eickhoff, Arjan Hillebrand, Tomas Knapen, Maartje C. de Jong, Serge O. Dumoulin

## Abstract

Vision is not instantaneous but evolves over time. However, simultaneously capturing the fine spatial details and the rapid temporal dynamics of visual processing remains a major challenge, resulting in a gap in our understanding of spatiotemporal dynamics. Here, we introduce a forward modeling technique that bridges high-spatial resolution fMRI with high-temporal resolution MEG, enabling us to non-invasively measure different levels of the visual hierarchy in humans and their involvement in visual processing with millisecond precision. Using fMRI, levels of the visual hierarchy were identified by measuring individuals’ population receptive fields and determining visual field maps. We predicted how much the activity patterns in each visual field map would contribute to brain responses measured with MEG. By comparing these predicted responses with the measured MEG responses, we assessed how much a given visual field map contributed to the measured MEG response, and, most importantly, when. We combined information from all MEG sensors and revealed a cortical processing hierarchy across visual field maps. We validated the method using cross-validations and demonstrated that the model generalized across MEG sensor types, stimulus shapes, and was robust to the number of visual field maps included in the model. We found that the primary visual cortex captured most of the variance in the MEG sensors and did so earlier in time than extrastriate regions. We also report a processing hierarchy across extra-striate visual field maps and clusters. We effectively combined the advantages of two very different neuroimaging techniques, opening avenues for answering research questions that require recordings with high spatiotemporal detail. By bridging traditionally separate areas of research, our approach helps close longstanding gaps in our understanding of brain function.

**Author Summary:** Vision doesn’t happen instantaneously, but unfolds over time. While we understand a lot about how the brain processes visual space, understanding how the brain processes information over both space and time is much more challenging. Vision happens incredibly fast, space and time are tightly linked, but current technology is limited in capturing these spatiotemporal dynamics.

Here, we developed a method that combines two non-invasive human brain imaging techniques using computational models: fMRI, which provides high spatial detail, and MEG, which captures millisecond-level timing. We use fMRI to model the detailed layout of the visual system, and predict how this spatial layout responds to specific visual stimuli and compare these predictions with MEG recordings. This allows us to pinpoint not just where visual processing happens, but also *when*.

Our results reveal a clear processing hierarchy: primary visual cortex responds first and explains most of the MEG signal, while higher-level visual areas activate later. We show that our method works across different stimuli, MEG sensor types, and brain areas, proving its robustness. This approach offers a new way to track how information flows through the living human brain, opening up new possibilities to study vision in both health and disease.

## Introduction

Vision is not instantaneous but evolves over time [1–3]. To rapidly interpret the world, the visual system relies on highly interconnected neuronal populations that form visual field maps that exchange information within hundreds of milliseconds [4–8]. This communication is dynamic, adapting to factors such as stimulus features, the current state of the visual system, and contextual influences, such as current task-demands [9–11]. Invasive recordings, mostly conducted in animal models, played a crucial role in uncovering the neural basis of visual processing. However, human visual processing may differ from animals in terms of visual system layout and differences in the amount of cognitive modulations from higher order regions of the brain [12].

To understand visual perception, we need to understand how information flows through the human brain. While invasive studies suggest a rough visual processing hierarchy, pinpointing the precise timing of neuronal responses across the visual cortex remains challenging in the healthy human brain. The challenge lies in the extraordinarily fast visual processing, the tight link between space and time, and the difficulty of measuring with high spatial and temporal resolution non-invasively in the living human brain.

Bridging the gap between spatial and temporal precision is essential to understand how visual information flows through the human brain. Functional magnetic resonance imaging (fMRI) provides high spatial resolution but lacks millisecond-scale timing. Magnetoencephalography (MEG) and other neurophysiological methods offer superior temporal resolution but struggle to pinpoint the exact cortical sources at millimeter resolution [3,13,14].

Here, we advanced pRF modeling approaches to estimate not only the contribution of individual visual field maps along the visual hierarchy, but also *when* they do so with millisecond precision. This method combines all MEG sensors to capture temporal information across cortical regions and builds on forward modeling approaches that combine the spatial precision of fMRI with the temporal resolution of MEG [15,16]. As a true model of neuronal activity, the model generalized over MEG sensor types and stimulus shapes, and was robust to the number of regressors used in the model, demonstrating the versatility and reliability of the method.

By using fMRI to predict pRFs at the cortical level, applying a forward model to project these into MEG measurement space, and evaluating response latencies, we provide detailed temporal activation windows of visual processing across the visual hierarchy. Specifically, primary visual cortex (V1) responded fastest, while extrastriate areas followed later on. Our modeling approach opens up new avenues to understand the differences in timing along the visual hierarchy under various circumstances, task-demands and clinical conditions.

## Results

To measure the time-courses of visual field maps along the visual hierarchy, we build a forward modeling approach, using population receptive field (pRF) models, to link high-spatial resolution fMRI and millisecond resolution MEG. First, we estimated population receptive fields using fMRI (Fig 1A Step 1; [17]) and defined visual field maps [18–21]. Second, we collected MEG responses to contrast-defined bar and circle shapes (Fig 1A Step 2). We used the pRFs models to provide a prediction of how they would respond to the same stimuli (Fig 1A Step 3). Next, we converted the pRF predictions to the MEG sensor-space, by applying masked gain matrices to the predictions that convert the visual field maps’ cortical responses to the sensor level (Fig 1A Step 4; [22,23]). The visual field maps’ sensor predictions were then compared to the measured responses in a cross-validated ridge regression [24], determining the contribution of visual field maps with millisecond resolution (Fig 1A Step 5).

**Fig 1.**
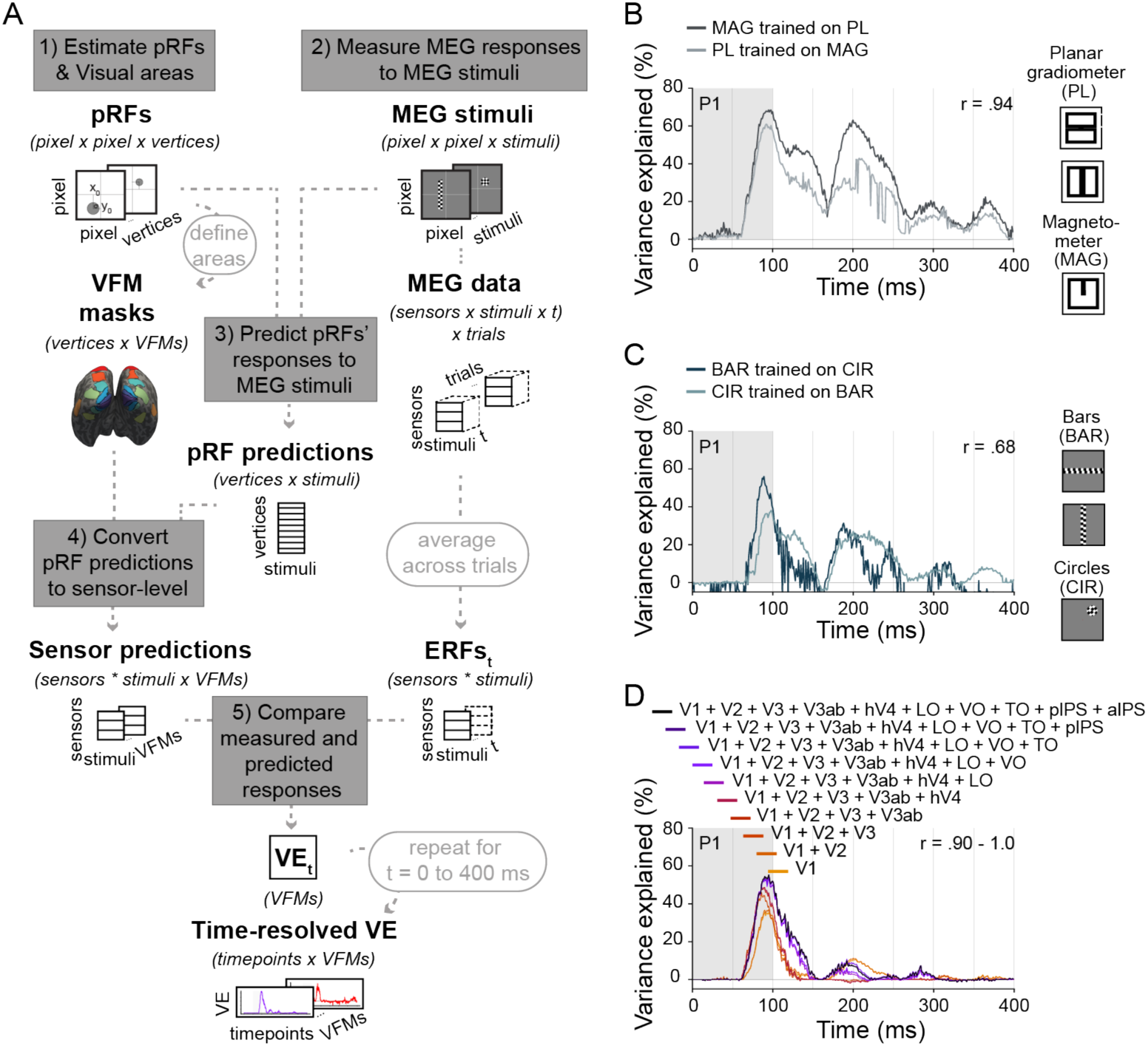
Overview modeling pipeline and model validation results. A. Modeling pipeline: *Step 1)* We measured each participant’s population receptive fields (pRFs) and defined ten visual field maps (VFMs) along the cortical surface. *Step 2)* MEG responses to bar and circle stimuli were recorded and averaged to event-related fields (ERFs). *Step 3)* pRF-based predictions were computed by matrix multiplying stimulus apertures with the pRFs. *Step 4)* The predictions were projected to MEG sensor space using a gain matrix. *Step 5)* A cross-validated ridge regression assessed how well VFM-based predictions explained the measured MEG response (variance explained; VE) at a given timepoint (t). **B**. **Sensor-type generalization**: Full model’s cross-validated variance explained for the model trained on one sensor type and tested on the left-out sensor type’s data for participant 1 (Fig S1A for all participants). The two time-courses were highly similar indicating our model generalized across sensor types (Pearson correlation coefficient: r(398)=0.94, p < .001). The gray shaded area indicates the period when the stimulus was shown. **C**. **Stimulus-type generalization**: Cross-validated variance explained for the model trained on one stimulus type and tested on the left-out stimulus type’s data (bars vs circles) for participant 1 (Fig S1B for all participants). Again the two time-courses were highly similar indicating that our model generalized across stimulus types (r(398) = .68, p < .001). **D**. **Robustness to number visual field maps**: We estimated V1’s cross-validated variance explained time-course including different numbers of extrastriate visual field maps for participant 1 (Fig S1C for all participants); i.e., ranging from including all ten visual field maps (black line), to only including V1 (yellow line). The time-courses were highly similar, indicating that V1’s time-course estimates were robust to the number of extrastriate visual field maps included in the model (r(398) = .90 – 1.0, p < .001).

### Model generalizes across sensors, stimuli and number of visual field maps

To assess the generalizability of our model, we trained the model on one of the sensor types, and tested the model fit on the left-out sensor type in all combinations, i.e., magnetometers versus planar gradiometers. The cross-validated estimated time-courses were highly correlated with the original time-courses, indicating high generalizability over sensor types (r(398) = .94, p < .001; r(398) = .96, p < .001; r(398) = .96, p < .001; r(398) = .93, p < .001; and r(398) = .98, p < .001 for participant 1-5, respectively; Fig 1B; Fig S1A for the other four participants).

In addition, in a similar procedure, we tested the generalizability of the models across stimulus shapes, i.e., bars vs circles. We used one stimulus type for training the model, and evaluated the model on the left-out stimulus type. The generalizability of the model was high across participants (r(398) = .68, p < .001; r(398) = .31, p < .001; r(398) = .32, p < .001; r(398) = .31, p < .001; and r(398) = .52, p < .001 for participants 1 to 5, respectively; Fig 1C for participant 1; Fig S1B for the other four participants).

We identified ten visual field maps and clusters, but we could have selected more or fewer. To assess the influence of the number of included visual field maps on V1’s estimated time-course, we calculated the fit using only V1 to explain the data, and then incrementally added all visual field maps up to including all ten areas in the model. We found high Pearson correlation coefficients between V1’s variance explained time-course that was obtained when only V1 was used in the model as compared to when the other visual field maps were included in the model. The minimum and maximum (i.e., the range) for each participant, was r(398) = .90 – 1.0, p < .001; r(398) = .91 - .98, p < .001; r(398) = .83 - .98, p < .001; r(398) = .97 - .99, p < .001; and r(398) = .99 – 1.0, p < .001 for participants 1-5, respectively, showing that V1’s fits across models with different numbers of predictors were stable for all participants (Fig 1D for participant 1; Fig S1C for the other four participants).

We also examined whether exclusion of V1 would alter the remaining visual field maps’ fits. We found that most visual field maps were stable when removing V1 from the model fit. For some visual field maps, namely those that had predictions that correlated strongly with V1’s predicted values, we observed changes in the variance explained time-course (Fig S2).

Together, the results suggest that our method is both generalizable and robust, allowing detailed investigation into the contribution of individual visual field maps in explaining visual responses, as well as their timing in doing so.

### V1 explains most variance in measured MEG responses

We calculated how much variance each visual field maps explained in the measured signal (Fig 1A Step 5). We found that V1 reached the highest variance explained across most participants (Fig 2B).

**Fig 2.**
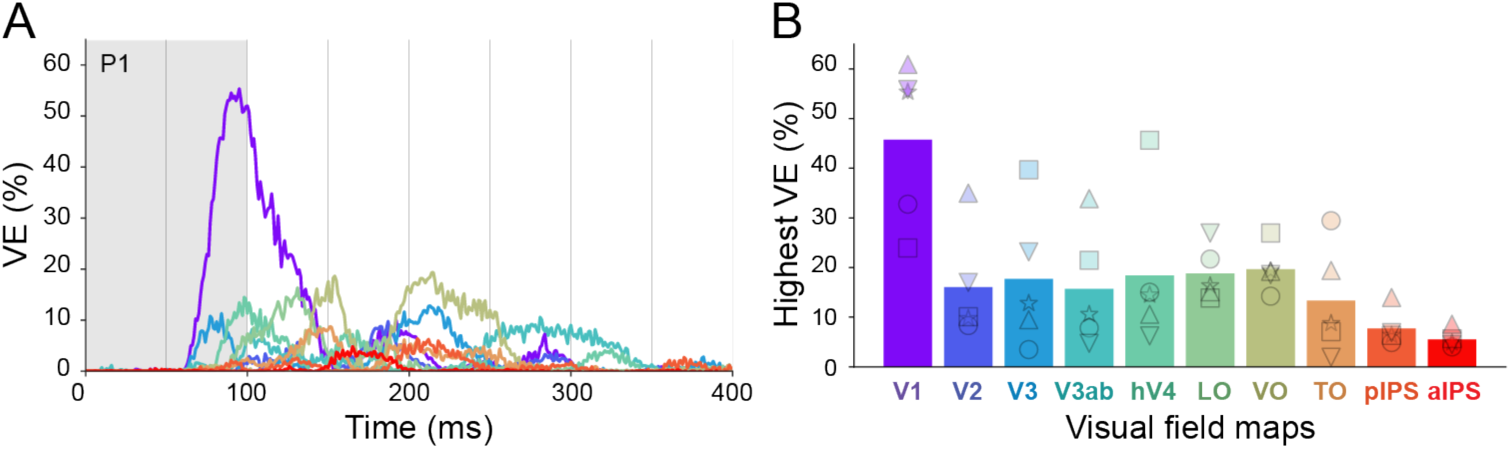
Visual field maps’ variance explained time-courses and maximal amplitude. **A**. Participant 1’s cross-validated variance explained (VE) time-courses for each visual field map from 0 to 400 ms after stimulus onset (color coded as in B; for other participants see Fig S3A). The gray shaded area indicates the period when the stimulus was shown. **B**. Visual field maps highest variance explained during the 400 ms time window. Bars indicate the mean over participants; individual markers each participant (star, circle, square, downward triangle, upward triangle for participant 1 to 5).

### Distinctive processing latencies across visual field maps

To quantify the timing of individual visual field maps, we identified the activity window in which they were contributing to the measured signal, by calculating when they reached 25, 50 and 75% of the normalized cumulative curve of the variance explained time-course (Fig S3B). This method avoids authors’ biases in setting thresholds and other parameters necessary for methods such as peak detection (Fig S4).

We found a trend of increasing latency along the visual hierarchy (from V1 to aIPS) across participants (Fig 3B). Across participants, the average timing of the 25th and 75th percentile of the cumulative VE time-course was 89 and 199 ms for V1, respectively, and 151 and 258 ms for the extrastriate regions.

**Fig 3.**
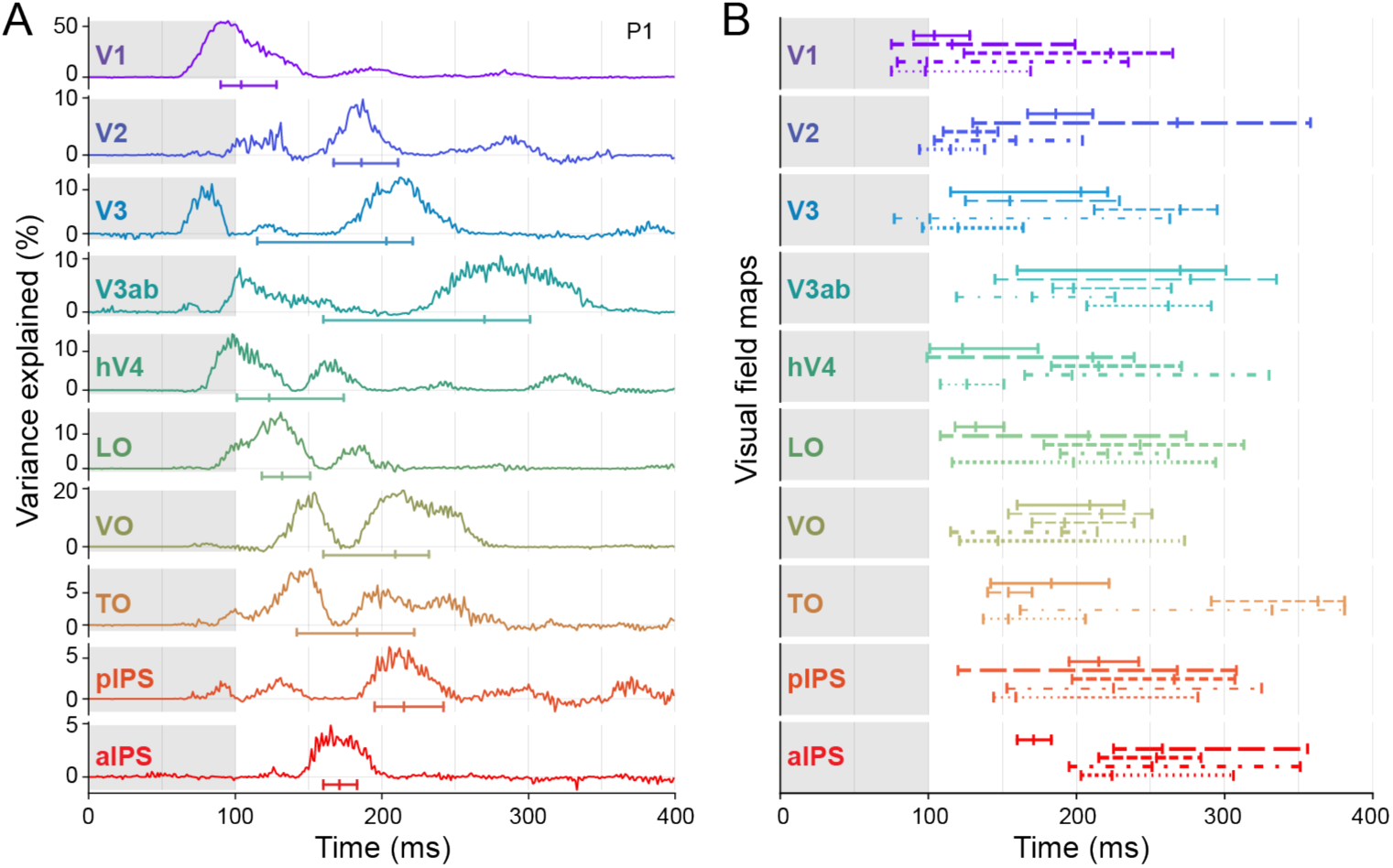
Variance explained time-courses and activation windows for different visual field maps. **A.** Cross-validated variance explained for different visual field maps (rows) for participant 1 for 0 to 400 ms after stimulus onset (see other participants in Fig S3A). The gray shaded area indicates the period when the stimulus was shown. The vertical ticks on the horizontal line below each variance explained time-course mark the time at which normalized cumulative variance explained reached 25, 50, and 75% of the total across the shown 400-ms time window. Note that the vertical axes are scaled differently for the different visual field maps. **B**. 25, 50, 75% of the normalized cumulative variance explained time-course (Fig S3B), for all five participants (P1: solid line, P2: long dashed line, P3: short dashed line, P4: dash-dotted line, P5: dotted line).

The forward modeling technique allowed us to investigate the contribution of individual visual field maps in explaining the measured MEG signal, and at which latency during the visual response they did so.

## Discussion

We showed that our new forward modeling technique captures when different regions of the visual hierarchy are active, and how much they contribute to the visual response (Fig 2). This was achieved by harnessing the spatial accuracy of fMRI, building computational models identifying visual processing regions of interest, and utilizing MEG to determine response timing, with millisecond resolution (Fig 3).

While multimodal neuroimaging using fMRI and MEG has been previously established (e.g. [25–27]) our approach builds on forward models of neuronal population responses [15,16] to precisely capture when specific visual field maps contribute to visual responses. We leverage pRF modeling [17], which describes a core property of brain function [17,28,29] and is build upon well-established neuronal properties (]30]; for review see [31,32]). pRF fMRI estimates correlate strongly with estimates from invasive neurophysiology in non-human primates [33], invasive human electrodes [34,35] and visual perception [36], and have proven stable, robust and reproducible [37–39], supporting their use as a computational framework for linking fMRI and MEG.

### Model generalizability and robustness

We validated our model by demonstrating generalizability across (i) sensor types, (ii) stimulus types and (iii) number of predictors (Fig 1B-D). Generalizability was somewhat lower across stimulus types than sensor types. This is expected as bar stimuli evoke more compressed responses and surround suppression than smaller circular stimuli [40–43]. These nonlinearities are not captured by our Gaussian model [17]. Advanced pRF models could improve the fitting [40,41,43,44], while aligning stimulus properties between fMRI and MEG experiments could further improve fitting results, e.g., including the circle shapes in the fMRI procedure. Nonetheless, our Gaussian model explained up to 66% of the response to a held-out stimulus, indicating it already provides a good estimate of the visual responses.

### Latency differences across visual field maps

We found V1 responded earlier than extrastriate regions, consistent with invasive electrophysiology showing hierarchical timing delays in both non-human primates and humans [2,45] and recent fMRI modeling [44]. This finding highlights the potential of our approach as a non-invasive tool for capturing high-temporal resolution brain dynamics, applicable for a wide range of human experiments.

The cumulative approach allowed us to investigate the timing of the visual field maps without making prior assumptions and offers an overview of the entire time window during which the individual areas are active and contribute to visual processing. We do not interpret the hierarchical delay as “one area follows the other”. There is an intricate interplay between regions along the hierarchy, with a general hierarchical trend of increasing latencies ([2,46]; see also [47]). Latencies may depend on the type of stimuli used [42,48–50], and other cognitive influences such as attention and task-demands [9,11]. Importantly, our approach enables future research to study how different tasks or cognitive states affect the contribution of different visual field maps, and to do so with ultra-high temporal resolution to detect any dynamic changes.

We found that V1 explained the most variance in most participants (Fig 2B and S3A-C). Based on the anatomy alone we would not expect V1 to dominate the signal: gain matrix values did not indicate higher sensitivity to V1 than to the other areas (Fig S5), and V1 did not contain more cortical surface points than the other visual field maps and clusters (Fig S6). Instead, we suggest that V1’s observed dominance in our data could be due to the fact that we used low-level contrast defined visual stimuli that strongly activate early visual regions and its highly organized parallel processing architecture [51–54]. It could be that more complex stimuli would activate higher order areas more strongly [55], resulting in a different distribution of visual areas contributions than observed here.

### Limitations

Our approach for measuring pRFs and the manual identification of visual field maps is time consuming for both the participant and the researcher. However, existing atlases may be used to reduce this burden [54,56,57]. Moreover, given the robustness of our results, we anticipate that fewer repetitions of stimuli could still yield accurate fits while requiring less time for the participant in the scanner.

Improved head models and signal space separation to correct for head movements could further improve the quality of the results [22,58,59]. Additionally, our approach is adaptable to other modalities. For instance, optically pumped magnetometers (OPMs), offer advantages over traditional MEG systems by providing better signal-to-noise ratios and accommodating head movements [60,61]. Our approach can extend to EEG as well, which is more widely accessible than cryogenic MEG / OPMs, by using appropriate individual head conductivity models [27,62–64].

## Conclusion

We non-invasively measured the contribution of individual areas along the visual hierarchy to visual responses in living humans. We did so with the same temporal resolution as invasive electrophysiological studies, by combining fMRI and MEG in a forward modeling approach. The computational framework is generalizable across experimental contexts and lays fundamental groundwork for investigating the dynamics of spatiotemporal processing both in health and disease.

## Method

### Experimental procedure and preprocessing

The full experimental procedure and preprocessing (Participants, Stimuli, Data acquisition and Data Preprocessing) is described in Eickhoff et al. [15].

In brief, five participants were measured in a separate fMRI and MEG session during which we presented contrast-defined stimuli, eliciting responses from similar neuronal populations across sessions. During both sessions, the participants were fixating on the center of the screen the entire time. Both fMRI and MEG data were minimally preprocessed before being fed into our analysis pipeline.

The functional MRI data was interpolated from voxel to vertex space to estimate pRFs on the cortical surface. The fMRI checkerboard stimuli (Fig 4A) and vertex data was used to estimate pRFs as described in Dumoulin & Wandell [17], and we subsequently predicted the pRFs responses to the MEG stimuli that were presented during the MEG session (Fig 4B and C).

**Fig 4.**
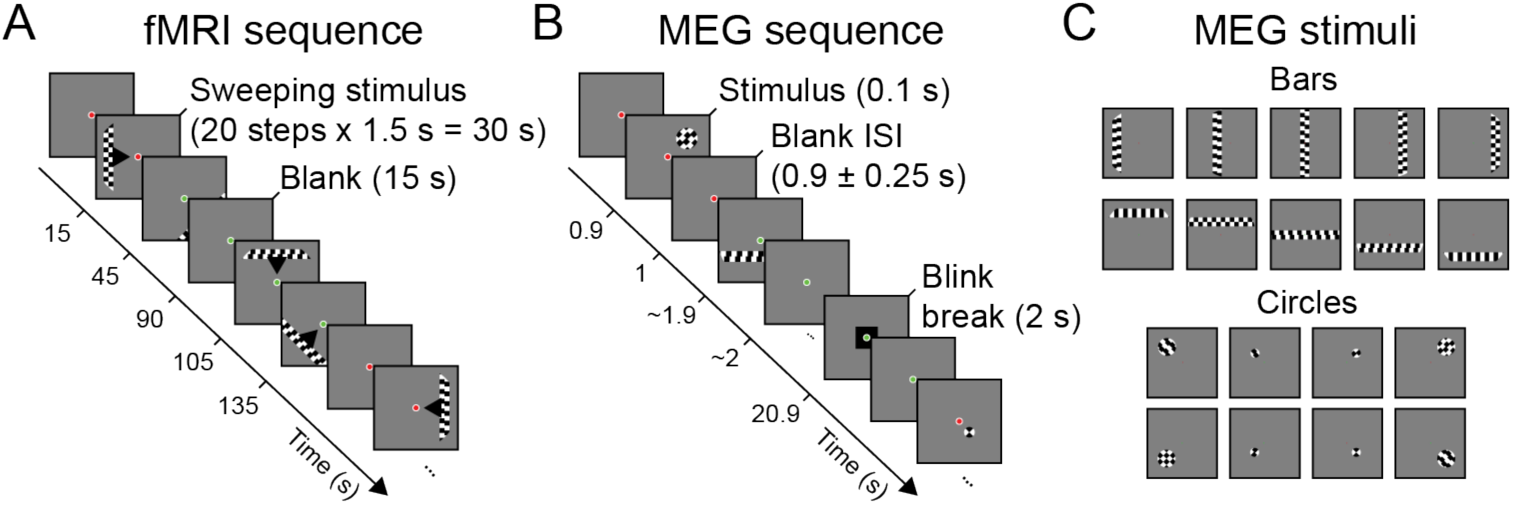
Stimulus sequence. **A**. A contrast-defined bar stimulus, interleaved by mean-luminance blanks, mapped the visual space in eight different directions while measuring fMRI. The stimulus sequence is a standard procedure to estimate pRFs (see [17]). **B**. Bar and circle shapes (shown in C) containing the same contrast-defined checkerboard were briefly (100 ms) shown in semi-random order, while measuring MEG. The stimulus trials were interleaved by mean-luminance blanks and blink periods. **C**. The eighteen MEG stimulus shapes, classified in two groups: ten *bars* (five vertical bars and five horizontal bars), and eight *circles*. The bars were either 0, 1.82 or 3.06 deg from screen center, the circles either 1.82 or 3.06 deg from center in the four visual quadrants, with a diameter of 1.25 and 3.06 deg respectively.

The MEG stimuli were eighteen stimulus shapes containing five vertical bars, five horizontal bars and eight circular shapes (Fig 4C), presented for 100 ms each in semi-random order. The MEG data was cut into time windows from 0 to 400 ms after stimulus onset, obtaining around 160 trials per stimulus.

The MEG data and the anatomical MRI scan were used to create the participants’ gain matrix. The gain matrix (MEG sensors x fMRI vertices) contains the weights of the vertices’ contribution to the signal measured with the MEG sensors. We estimated the gain matrix with the overlapping spheres (OS; [22]) model, and constrained the dipoles to be perpendicular to the cortical surface.

The fMRI-based pRFs and gain matrix were used, together with the MEG data, to perform our forward modeling approach.

### Forward modeling approach

To estimate spatiotemporal responses to visual stimuli in the visual system with millisecond resolution, we applied a forward model that converts visual field maps-specific predictions to MEG sensor level, allowing us to estimate how much a given visual field maps contributes to the measured MEG response. In brief, fMRI was used to estimate participants’ pRFs and define visual field maps (Fig 1A Step 1 and Fig 5) and MEG sensor responses were collected in response to MEG stimuli (Fig 4C and Fig 1A Step 2). Next, we predicted how the pRFs would respond to the same MEG stimuli (Fig 4C and Fig 1A Step 3). Then, we confined the predictions to one of the ten defined visual field maps (VFM). For that, the predictions were converted to sensor-level by applying VFM-masked gain matrices (Fig 1A Step 4), and we computed how much each of the predicted responses explained the measured MEG data in a cross-validated manner (Fig 1A Step 5). To demonstrate the generalizability and reliability of the approach, we performed three method validation tests, namely 1) testing whether the model generalized over MEG sensor types (magnetometers vs planar gradiometers), and 2) stimulus shapes (bars vs circles), and 3) testing whether the model results were robust to the number of predictors used in the model. The analysis steps are explained in more detail below.

**Fig 5.**
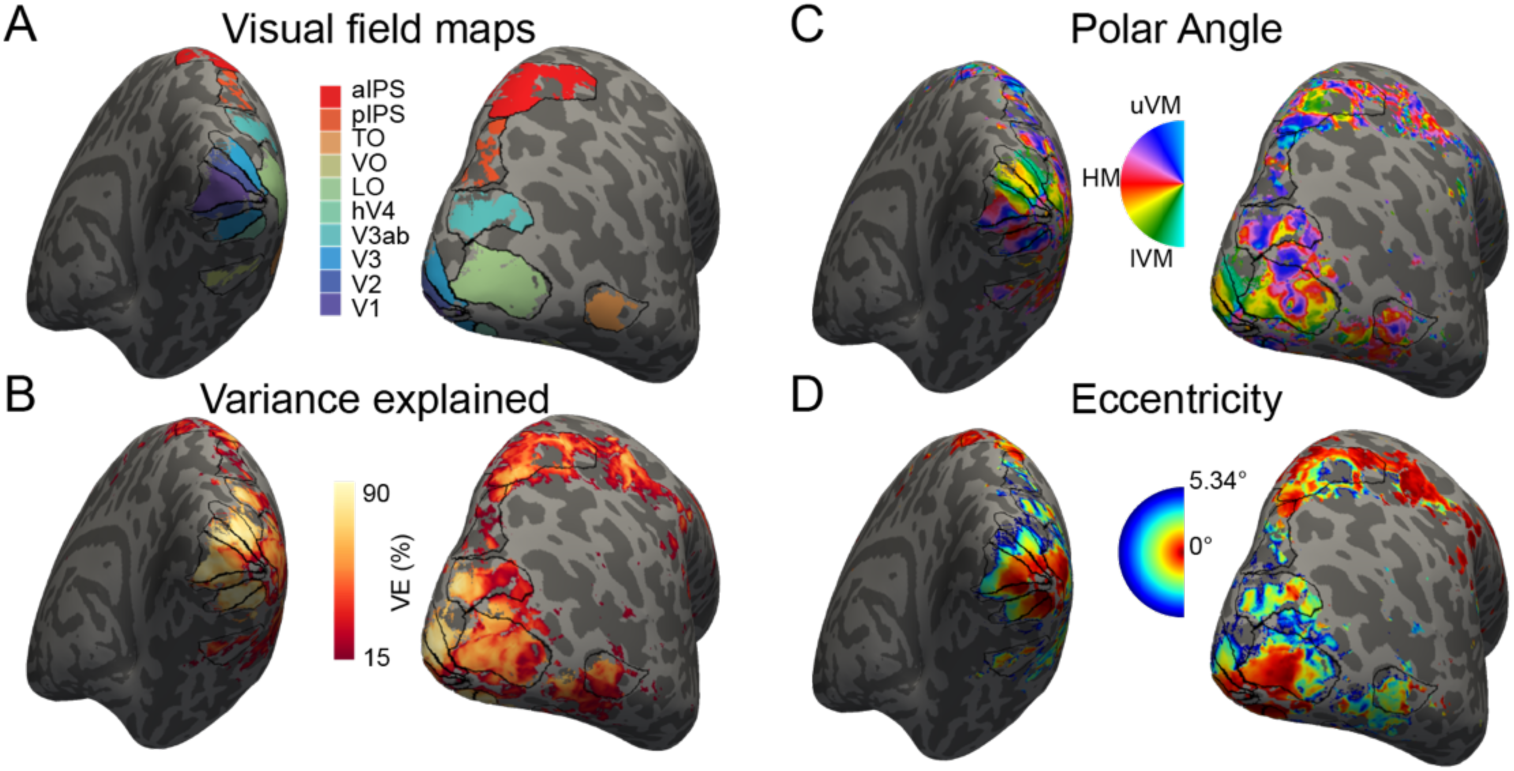
Visual field maps. The left column in each panel shows a medial posterior view of participant 1’s cortical surface. The right column a lateral posterior view. Only vertices included in the analysis are color-coded. **A**. Visual field maps that were defined by hand, based on the visual field maps shown in row two to four. **B**. How much variance a given pRF explained in the fMRI data. **C**. The polar angle in visual space associated with a given pRF, ranging from the upper vertical meridian (uVM), through the horizontal meridian (HM) to the lower vertical meridian (lVM). **D**. The eccentricities in visual space of the pRFs; 5.34 deg was the maximum eccentricity at which the stimuli were presented.

#### Step 1: Estimate pRFs and visual field maps

To estimate the participants pRFs, we measured fMRI while participants viewed contrast-defined bar stimuli that moved across the visual field (Fig 4A). For each cortical location (called ‘vertex’), a pRF was estimated as previously described in Dumoulin & Wandell [17]. A brief overview follows:

The accumulated receptive fields of a population of neighboring neurons can be approached as a 2D Gaussian in visual space. A range of these 2D Gaussian models with different visual locations were constructed, and their responses to the bar stimuli predicted. The predicted time-courses are the result of matrix multiplication of the 2D binarized stimulus and the 2D Gaussian. The predicted time-courses are then fitted to the measured time-courses to the same stimuli for a given cortical location, and the Gaussian that fits best was selected as the estimated population receptive field (pRF) for that vertex. The pRF thus described the part of the visual field to which a cortical location responds the most. For the following analysis steps, we only used vertices with pRFs that had fits with variance explained (VE) of at least 15%. We also excluded vertices that were located under a vein, and where the pRF eccentricity was located outside the visual space in which the stimuli were presented.

We then defined visual field maps for each participant (Fig 5A), based on the measured visual field maps (Fig 5B-D). For that we examined anatomical landmarks and polar angle reversals and eccentricity values. Based on the measured visual field maps we identified ten visual field maps here: V1, V2, V3, V3ab, hV4, LO, VO, TO, posterior IPS (pIPS) and anterior IPS (aIPS). Note that we could have merged certain regions together (for example, posterior and anterior IPS), or separate them even further. However, for the purpose of this method, the current level of specificity matched other pRF modeling papers [40,45]. Furthermore, we addressed the question regarding the effect of the choice of the number of visual field maps as regressors in our Method validation section.

#### Step 2: Measure MEG responses to MEG stimuli

We measured MEG data with 306 sensors, while participants viewed different bar and circle stimuli. The stimuli were eighteen different stimuli in total (Fig 4C), each repeated on average 160 times in semi-random order (Fig 4B). Each stimulus was presented for 100 ms to evoke strong event-related responses, and the responses were cut into trials from 0 to 400 ms after stimulus onset, obtaining a matrix of (sensors x stimuli x timepoints x trials; Fig 1A Step 2).

#### Step 3: Predict pRFs’ responses to MEG stimuli

We next predicted how the pRFs on the participants’ cortical surface respond to each of the MEG stimuli shown. The predicted response is the spatial overlap between the 2D binarized stimulus and the 2D pRF models, calculated as matrix multiplication. This step resulted in a matrix of (vertices x stimuli), containing each pRF’s response to each of the eighteen stimuli (Fig 1A Step 3).

#### Step 4: Convert visual field maps’ pRF responses to MEG sensor-level

To identify whether the pRFs measured on the cortical surface can explain the MEG data, the pRFs responses need to be converted to the MEG measurement space. That is, while the pRFs predictions live in cortical space, the MEG data was measured at the sensor-level. The gain matrix determines how much each vertex contributes to a given sensor, hence multiplying the pRF predictions with the gain matrix converts the predicted cortical values to sensor-level: matrix-multiplying the gain matrix (sensors x vertices) with the cortical pRF predictions (vertices x stimuli), scales each vertex’ response by the appropriate amount for each MEG sensor, resulting in a matrix of (sensors x stimuli).

We do not expect that each visual field map contributes the same amount at a given timepoint during the visual response. Instead, we expect that distinct visual regions would reveal different response time-courses. Early visual field maps should respond early and receive feedback later in time. While higher order areas respond later as the signal needs time to travel up the cortical hierarchy first. We thus created visual field map-specific predictions by letting only the vertices belonging to a given area weigh into the predicted sensor-level responses. More specifically, we created ten visual field maps-masked gain matrices (each sensors x vertices), in which the weights of all vertices outside a specific area were set to zero, and then multiplied the visual field maps masks with the gain matrix for the participant. Thus, only the given visual field map’s vertices now contributed to the sensor-level prediction. By matrix-multiplying each masked gain matrix with the vertex’ responses, we thus obtained ten visual field maps-specific sensor predictions (Fig 1A Step 4), each with shape (sensors x stimuli).

#### Step 5: Compare measured and predicted responses

Lastly, we calculated how well each visual field maps-specific prediction fitted the MEG data using cross-validated ridge regression, for each of the timepoints of the visual response after stimulus onset. We opted for ridge regression since it allows for penalization of beta weights of correlated visual field maps predictors.

For the cross-validation procedure, we obtained four sets of average responses (referred to as ‘event related field’ (ERF) responses) from the MEG data (Fig 1A Step 2 bottom). The averaging gets rid of unrelated noise while preserving stimulus-related responses in the signal. The averages were drawn from four random splits of trials per stimuli, resulting in four sets of (sensors x stimuli x timepoints). One of the sets was left out, and was used later as independent evaluation of how well the model predictions fit never-seen data (test set). The three other sets were used in a three-fold cross-validation method to find the parameters that resulted in the best fit between the ten visual field maps predictors and the measured data.

Before the data matrices were fed into the ridge regression, the datapoints across all sensor and stimuli were concatenated to a big matrix of (sensors*stimuli) for both predictions and measured data. The visual field maps predictions were thus a matrix of (sensors*stimuli) x visual field maps, and the measured data for a specific timepoint *t* was (sensors*stimuli).

To perform ridge regression at a given timepoint, we implemented the *fracridge* toolbox in a cross-validated manner [24]. This procedure ensures the appropriate level of regularization is applied by reparametrizing the regression based on the a given ratio (gamma) between the regularized and unregularized L2-norm coefficients. For a range of ratios (gammas) from 0.1 to 1 in steps of 0.1 the toolbox tested which ratio and accompanying regularization parameter (alpha) resulted in the best fit in a three-fold cross-validated manner: For a given gamma, one of the three datasets was left out for testing. The other two datasets were used to obtain the alpha parameter and beta values for each predictor. The betas were fixed and applied to the predictions to test on the left-out third set, obtaining the goodness of fit for that particular fold. This was repeated for the other two datasets. The three parameter sets resulting from this were then averaged, and the whole procedure repeated for the other gamma ratios. The gamma value and belonging parameters resulting in the highest goodness of fit were chosen, and applied to the predictions of the fourth, never-seen, test set, and the goodness of fit (r^2^) was computed. This goodness of fit value represents how well the full model (all predictors/visual field maps) explained the measured data at a given timepoint. We repeated this for all timepoints (0 to 400 ms) (Fig 1A Step 5). In the present dataset, regularization was low when visual signal was present (Fig S7).

Finally, since we were interested in the individual contribution of the predictors (visual field maps; VFM) in explaining the measured data, we calculated how much variance (VE) each visual field map explained in the data as follows:

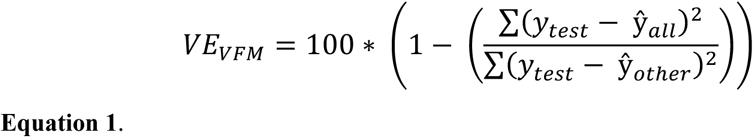

where *y_test_* contains ERF values (sensors*stimuli,) at timepoint t of the test set, 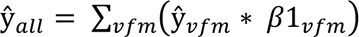, and 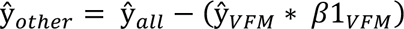.

To quantify the latencies of the individual visual field maps’ responses, we used a cumulative approach that takes the total history of the variance explained time-course into account and thus characterizes the entire ‘activation window’ of an area. This approach avoids assumptions and experimenters choices on parameters that would be necessary for other approaches such as peak detection.

For the cumulative approach, we calculated the normalized cumulative VE curves based on each variance explained time-course (Fig S3B). For that, we set timepoints with negative variance explained to zero and calculated each timepoint’s cumulative VE value as the sum over its current value and all past values up to that latency and the cumulative time-course was normalized to values between 0 and 1. We then examined the timepoints at which the normalized cumulative VE reached 25%, 50% and 75%.

## Method validation

We performed three validation tests. We examined, 1) how well the model generalized across the different sensor types of the MEG scanner (magnetometer vs planar gradiometers), 2) how well it generalized across the stimulus types used (bars vs circles), and 3) how robust the results were to the number of predictors (visual field maps) included in the model.

### Sensor type validation

The MEG system used to measure the data holds two sensor types, which have different signal sensitivity profiles [65]. We wanted to test whether the model generalizes across these two sensor types. Instead of using all sensors for cross-validated fitting for a given train-test fold, we only used one sensor type for model training. The model performance was then evaluated on the left-out sensor type. We left out each sensor type once, resulting in two variance explained time-courses. Within each train-test combination, we used the same cross-validated three-fold procedure, i.e., the concatenated data of the ‘train’ sensor types was split into three sets to find the optimal regularization. Below, we report the final goodness of fit of the full model, calculated as:

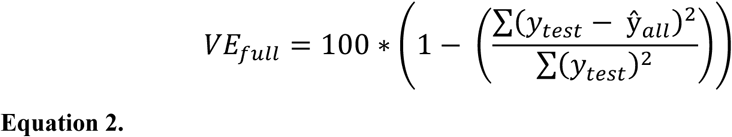

where *y_test_* contains ERF values (sensors*stimuli,) at timepoint t of the test set and 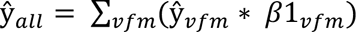. We report the Pearson correlation coefficient between the train-test results to quantify the stability across sensor type fits.

### Stimulus type validation

We tested the model’s generalizability across the responses to the different stimulus-shapes used. The two stimulus types were the ten bars and the eight circular stimuli (Fig 4C). Again, we used only one of the stimulus shapes for training of the model, and used the left-out stimulus type to evaluate the fit. We used the same cross-validated three-fold ridge regression procedure for each combination of train-test stimulus types, and again reported the final goodness of fit of the full model (Equation 2) for each fold and the correlation coefficient across these train-test folds.

### Robustness to number visual field maps

The robustness of visual-fits to the number of predictors (i.e., the visual field maps) in the model was examined by evaluating models with different numbers of predictors. We focused on the V1 time-course and its stability as it resulted in the best variance explained across most participants (Fig 2). For this, we used all stimuli and sensors’ data; however, the predictor matrices now originated from different numbers of visual field maps. That is, first we used only the predicted values from V1, then incrementally added each visual field map until the model contained the full ten visual field maps. We performed the same three-fold cross-validation procedure as described above, leaving out a fourth, independent set of data to test the model. We reported the V1 prediction fit (as calculated in Equation 1), for each of the models tested and to quantify the stability, and reported the correlation coefficient between the V1-only fit to the other nine model fits. The effect of removing V1 (the main predictor; Fig 2) on the stability of the fits of the other visual field maps was examined by excluding V1 from the model fits. The results can be found in Fig S2.

## Supporting Information

**Fig S1.**
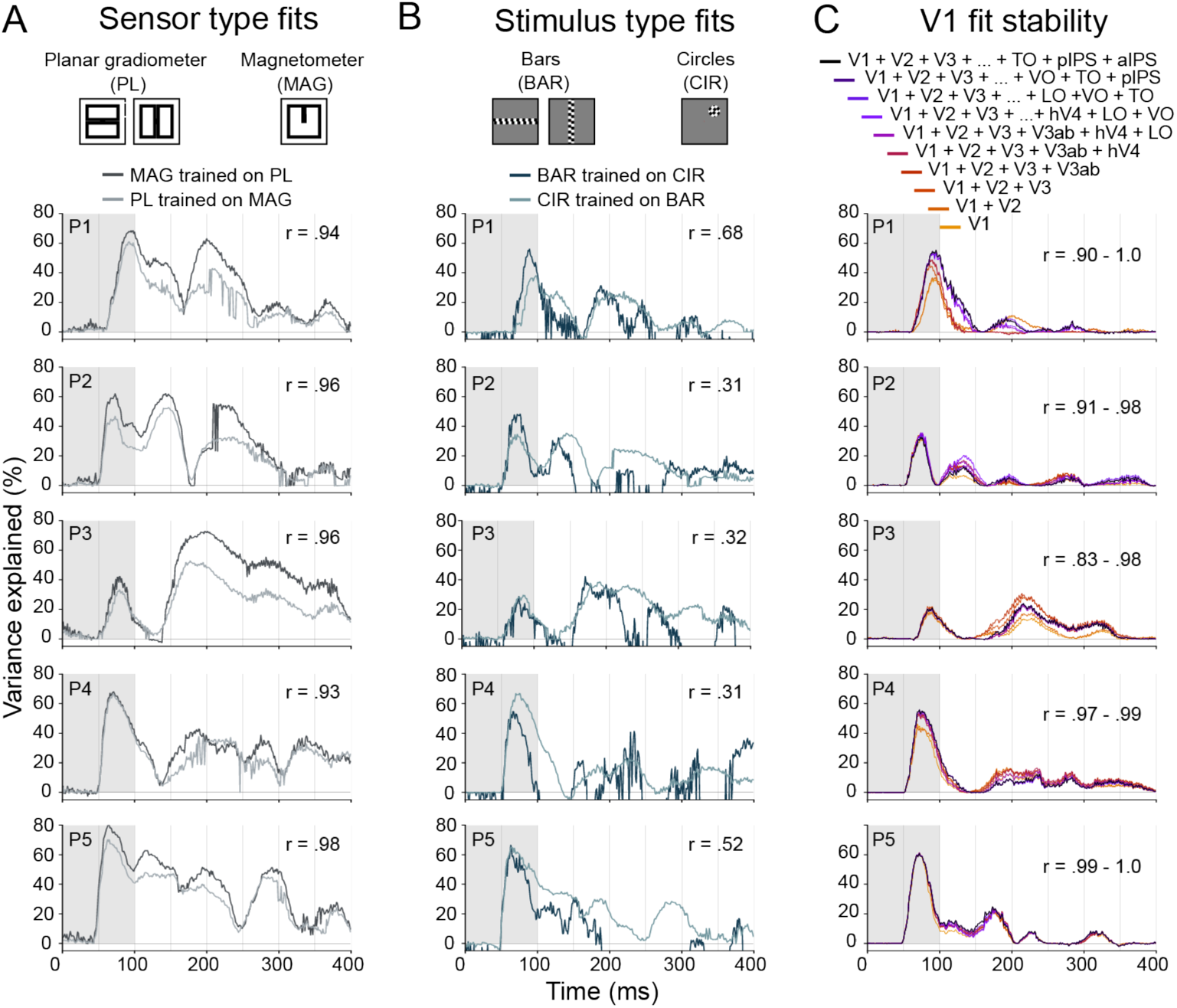
Method validation results for all participants. **A**. Cross-validated variance explained (*Methods* Equation 2) for the model trained on one sensor type and tested on the left-out sensor type’s data for all participants (rows). The Pearson correlation coefficient of the two time-courses (shown in upper right corner) was high, ranging from .93 to .98 across participants, indicating our model generalized across sensor types. **B**. Cross-validated variance explained for the model trained on one stimulus type and tested on the left-out stimulus type’s data (bars vs circles) for all participants (rows). **D**. V1’s cross-validated variance explained time-course (*Methods* Equation 1) for models with different numbers of visual field maps for all participants (rows); ranging from including all ten visual field maps (black line), to only including V1 (yellow line). Presented r range (upper right corner) reflects the relationship of the V1-only model fit to the other nine model fits. The high r (ranging from 0.83 to 1.00) indicates that V1’s fit was stable across different numbers of visual field maps included in the model.

**Fig S2.**
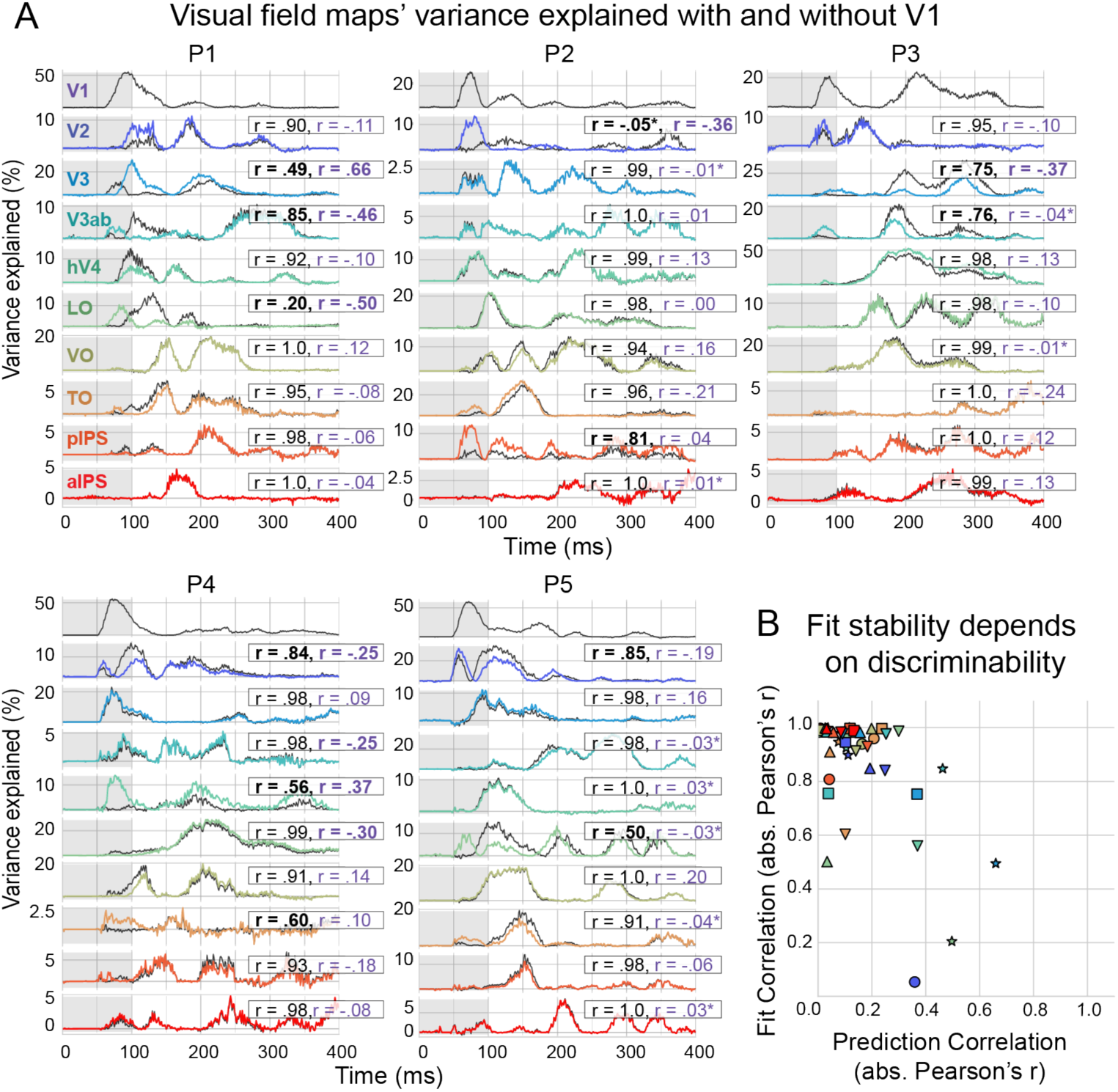
Visual field maps’ variance explained fitted with and without V1 regressor. **A**. For each participant (P1 to P5) and visual field maps (rows): cross-validated variance explained time-course resulting from two model fits: when V1 was included in the model (black lines; same model fit as in main analysis), and when V1 was excluded from the model fit (colored lines). In each panel the correlation between the two fits (“fit correlations”) are indicated in black. All fit correlations had a p-value of <.001 (N = 398), except for V2 in participant 2 (\**p* = .28. For most visual field maps across participants these correlations were high, indicating that the visual field maps’ fits were stable and independent of V1’s inclusion in the model. Some visual field maps’ fits changed, which is reflected in lower correlation values; these are highlighted in bold (r ≤ .85 or ≥ -.85). The low fit correlations often occurred together with high “prediction correlations” (indicated in purple, and highlighted in bold if r ≥ .25 or ≤ -.25); which are the correlations between the respective visual field maps’ predicted values and V1’s predicted values, indicating that the regressors cannot be distinguished well by the model. All prediction correlations had a p-value of < .001 (= 398), except those with stars; deviating p-values in order of participants, top to bottom are: P2 V3: *p* = .349, P2 V3ab: *p* = .531, P2 pIPS: *p* = .002, P2 aIPS: *p* = .513; P3 V3ab: *p* = .005, P3 VO: *p* = .288; P5 V3ab: *p* = .019, P5 hV4: *p* = .047, P5 LO: *p* = .013, P5 TO: *p* = .001, P5 aIPS: *p* = .019. **B**. Summary plot of the Fit and Prediction correlations plots. Each color corresponds to the visual field maps; participants 1 to 5 had markers: star, circle, square, downward triangle, upward triangle, respectively. Most Fit correlations were high (above 0.85), indicating stable fits independent of V1 in- or exclusion. Lower Fit correlations were related to higher Prediction correlations.

**Fig S3.**
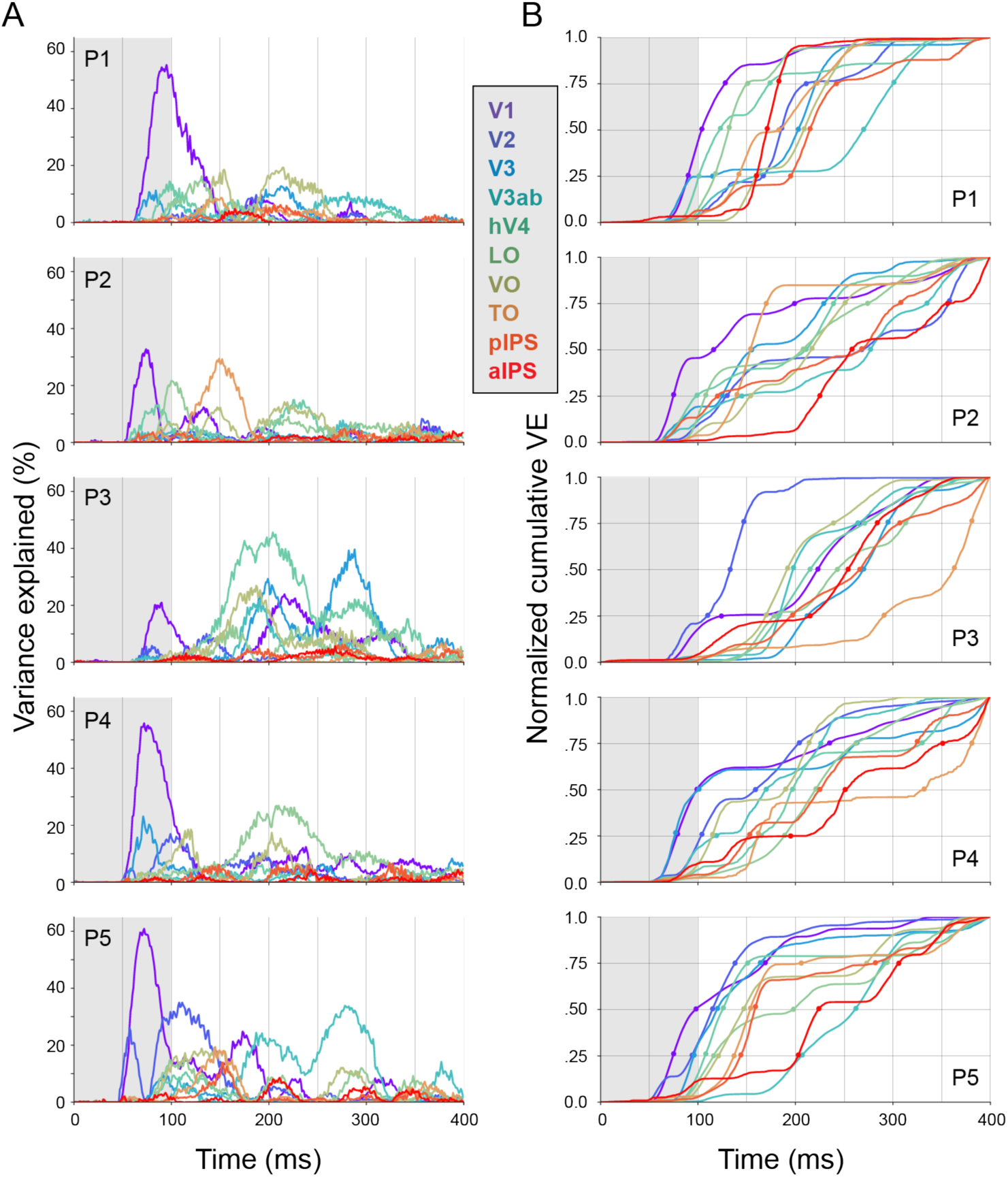
**A**. Cross-validated variance explained time-courses of each visual field map (see middle inset for color code) for all participants (rows P1-P5) from 0 to 400 ms after stimulus onset. The gray shaded area indicates the period when the stimulus was shown. **B**. Normalized cumulative variance explained (VE) for each visual field map and participant.

**Fig S4.**
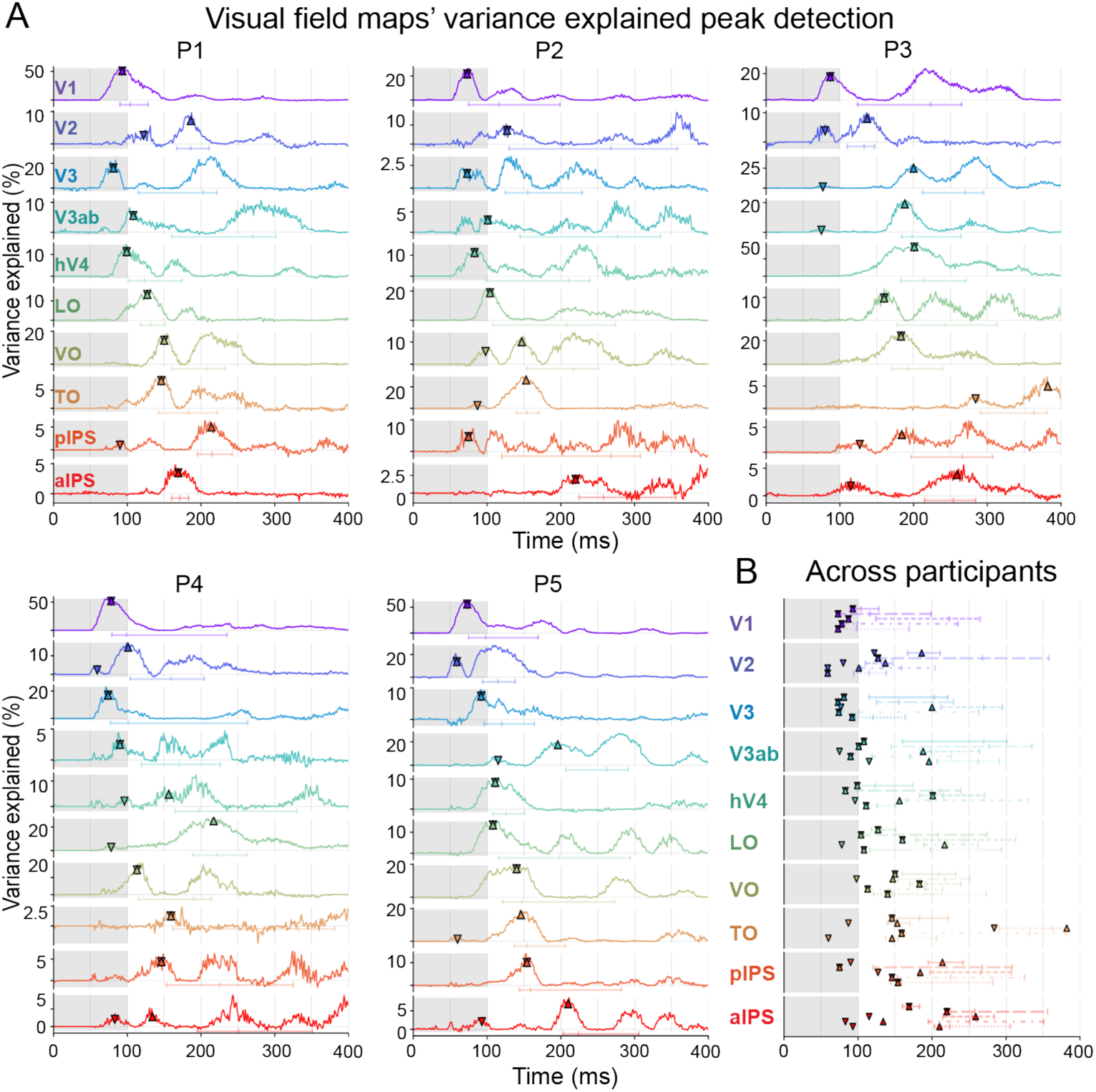
Using peak detection for latency quantification. **A**. For each participant (P1 to P5) and visual field maps (rows), we show the peaks found for two peak detection methods (markers ‘v’ and ‘^’ for method 1 and 2, respectively), on top of the cross-validated variance explained (VE) time-courses for each visual field map (note the different vertical scales). For both methods we smoothed the VE time-courses with a 20 ms moving window, and identified local maxima with a width of at least 10 ms, but we applied different thresholds: for method 1 we considered all peaks with variance explained above 0; for method 2 we only considered peaks with a variance explained of at least 50% of the maximum variance explained. For comparison, below each VE time-course are the latency window results found in our main analysis; vertical ticks mark the latencies at which normalized cumulative variance explained reached 25, 50 and 75% of the total across the shown 400-ms time-window. **B**. Peaks and cumulative time windows for all subjects (P1 to P5 from top to bottom) for each visual field map. For method 1, the average first peak detected was 81 and 112 ms for V1 and extrastriate regions, respectively; for method 2, the average peak was 81 for V1 and 141 ms for extrastriate regions. Both peak detection approaches resulted in earlier latencies than the activation time window determined by the cumulative approach with an average 25% percentile onset of 89 and 151 ms for V1 and extrastriate regions, respectively.

**Fig S5.**
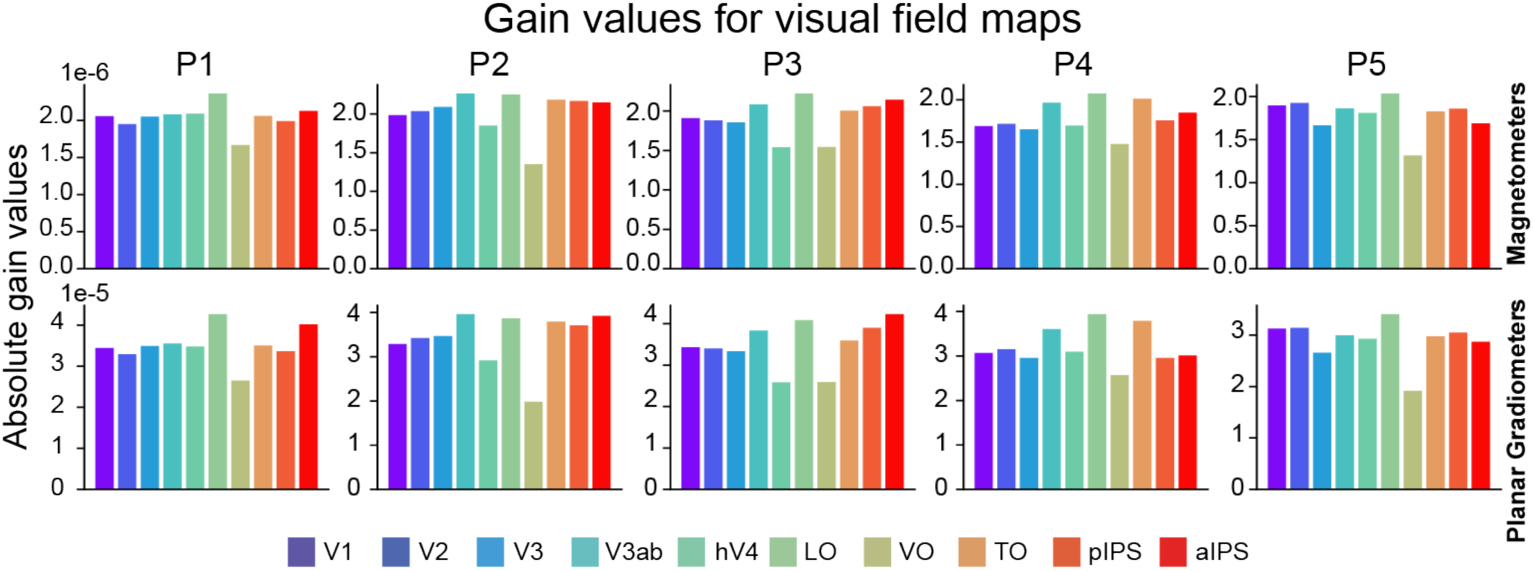
Gain values for visual field maps. Absolute gain values averaged over 102 magnetometers and 204 planar gradiometers, for each visual field map and participant separately.

**Fig S6.**
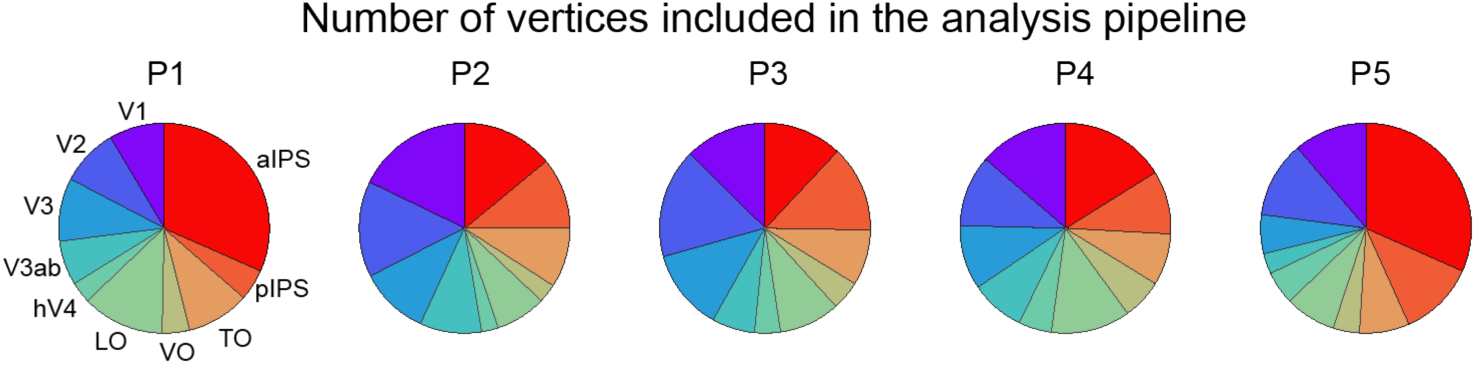
Number of vertices per visual field map included in the analysis for each of the five participants.

**Fig S7.**
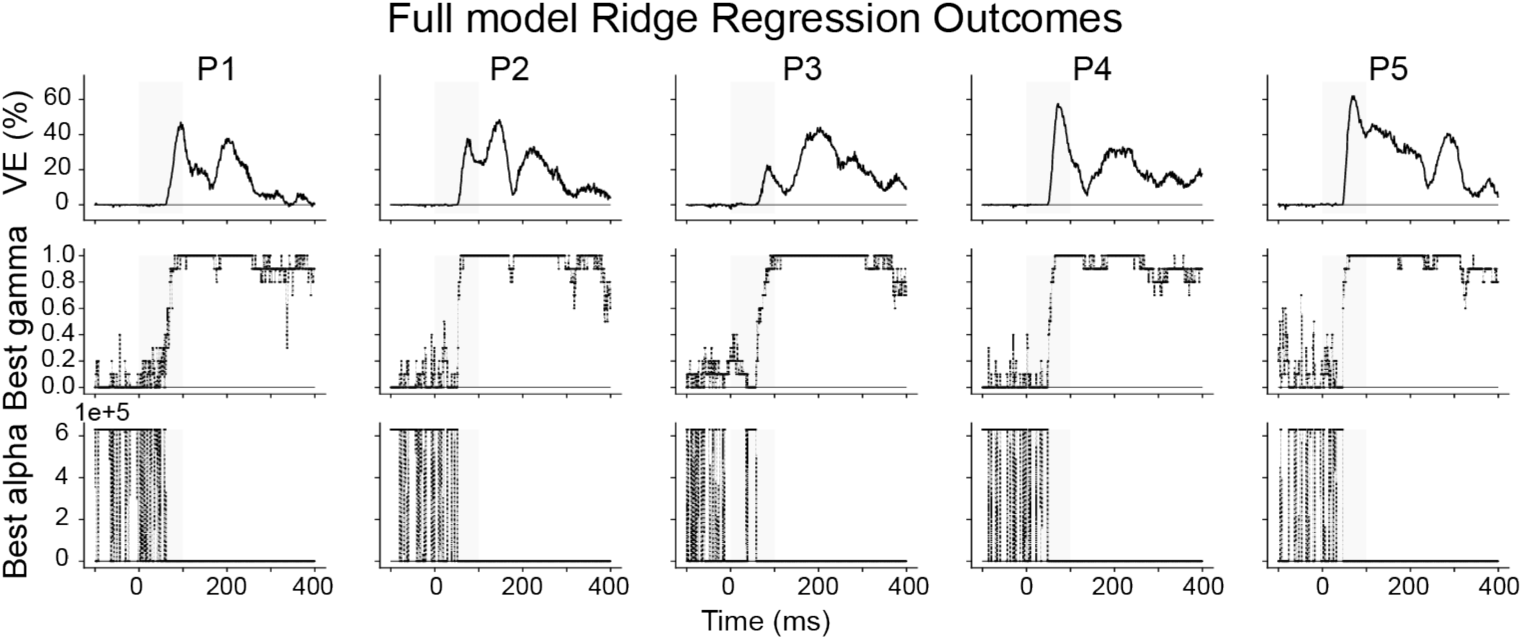
Full model ridge regression cross-validation outcomes. For all five participants (columns). *Top row*: Full model variance explained (VE) time-courses for all participants calculated as in Equation 2. This time-course shows how much all visual field maps together explain the measured MEG data. *Middle row*: Best average gamma (ratio) found for each timepoint with the three-fold cross-validation procedure. A value of 1 corresponds to no difference between the regularized and non-regularized outcomes. High gamma values correspond to low alpha (regularization parameter) values, as shown in the *bottom row*: High alpha values indicate high regularization (high punishment of beta values, for example if predictions are correlated). An alpha value of 0 corresponds to no regularization, so ordinary least squares. These results indicate that when the signal is present, generally no regularization was required.

